# A quantitative framework for predicting odor intensity across molecule and mixtures

**DOI:** 10.1101/2025.08.08.668954

**Authors:** Robert Pellegrino, Khristina Samoilova, Yusuke Ihara, Matthew Andres, Vijay Singh, Richard C. Gerkin, Alexei Koulakov, Joel D. Mainland

**Affiliations:** Monell Chemical Senses Center, Philadelphia, Pennsylvania, USA; Cold Spring Harbor Laboratory, Cold Spring Harbor, NY, USA; Institute of Food Sciences and Technologies, Ajinomoto Co., Inc., Kawasaki, Kanagawa 210-8681, Japan; Department of Physics and Astronomy, Haverford College, Haverford, PA, USA; School of Life Sciences and School of Mathematical and Statistical Sciences, Arizona State University, Tempe, AZ, USA; Osmo Labs, PBC, Cambridge, MA, USA; Department of Neuroscience, University of Pennsylvania, Philadelphia, PA 19104, USA

**Keywords:** olfaction, biophysics, human perception, psychophysics, intensity

## Abstract

In vision and hearing, standardized units such as lumens (for brightness) and decibels (for loudness) allow consistent quantification of stimulus intensity, enabling precise control of sensory experiences. Olfaction, by contrast, currently lacks a robust quantitative framework linking physical stimulus properties directly to perceived odor intensity, complicating efforts to accurately characterize and manipulate aromas. To bridge this gap, we used a precisely controlled odor delivery system combined with deep learning models to predict the intensity of both single molecules and mixtures from physical properties. These models allowed us to develop an automated, quantitative method that accurately identifies which volatile components meaningfully contribute to aroma perception, overcoming the limitations of traditional heuristic approaches such as odor activity values and demonstrating practical utility in complex naturalistic odors.

## Introduction

The number of photons or the amplitude of air pressure variations, respectively, can determine the magnitude of visual or auditory stimuli, thus providing a simple metric for representing quantity using a single physical property. In these sensory modalities, an understanding of intensity was necessary to elucidate how the nervous system encodes stimuli and how individual components contribute to a mixture. In contrast, the concentration of an odorant alone is a poor predictor of the perceived intensity, with detection thresholds and maximum intensities varying widely across odorants. To develop the olfactory equivalent of noise-canceling headphones, high-fidelity audio, or digital videos we must first be able to predict the perceived intensity of an odor.

Previous research on olfactory modeling has established transport-based boundaries for whether a molecule is odorous (Mayhew et al. 2022), predicted detection thresholds (Abraham et al. 2012), and predicted intensity at specific concentrations (Edwards and Jurs 1989; Keller et al. 2017). However, the field still lacks a quantitative axis for perceived odor strength, especially in mixtures, where intensity dictates which components dominate perception. Human and rodent psychophysical studies demonstrate that intensity does not map linearly to concentration, either within or across odorants (Sirotin et al. 2015; Wakayama et al. 2019; Wojcik and Sirotin 2014). Consequently, existing quantitative models cannot extrapolate the intensity at different concentrations. The relationship between an odor’s concentration and intensity is well described by a three-parameter Hill equation (Chastrette et al. 1998), yet there is no comprehensive dataset describing Hill parameters across odorants, or models predicting these parameters from molecular structure. Moreover, traditional psychophysical models for mixtures typically rely on single-concentration intensity measurements of individual odorants (Cain et al. 1995), despite evidence suggesting that full concentration-intensity curves are essential for accurate predictions of mixture intensities (Singh et al. 2019). This gap is especially significant given the complexity of natural odors, where intensity, rather than concentration, better predicts shifts in odor character for specific odorants (Gross-Isseroff and Lancet 1988; Laing et al. 2003), and more accurately weights the contribution of odorants in complex mixtures (Ravia et al. 2020).

Several critical barriers have hindered progress toward reliable intensity prediction in olfaction. First, there is a shortage of perceptual data, particularly for mixtures, limiting the accuracy and applicability of existing models (Gerkin 2021). Second, precise stimulus control has been difficult, with most studies reporting liquid-phase concentrations despite the need for accurate air-phase (headspace) concentration measurements due to variability in ambient dilution during a sniff, solvent interactions (Jennings et al. 2023), and odorant binding to delivery systems (Gorur-Shandilya et al. 2019). Finally, two odorants with similar intensities but disparate binding affinities will have very different effects on a mixture’s intensity, necessitating data collection for a broader set of concentrations. Addressing these barriers, we developed a robust, quantitative modeling approach using an extensive psychophysical dataset comprising full concentration-intensity profiles for individual odorants (62 molecules at seven or more concentrations) and comprehensive mixture data. This method substantially improves intensity prediction, thereby enhancing stimulus control and advancing olfactory research toward the precision achieved in other sensory modalities.

Here we demonstrate that a comprehensive, high-quality psychophysical dataset of mixture intensities (N = 260), alongside concentration-intensity functions for individual molecules (62 molecules at seven or more concentrations, totaling 434 data points), enables the prediction of odorant and mixture intensity based on molecular structure. Notably, biologically inspired models excelled in predicting odor mixture intensity, and molecular features crucial for transporting an odorant from its source through mucosa to the odorant binding site were predictive of monomolecular intensities. We extend the application of these intensity models to predicting odor quality, revealing that they enhance performance compared to leading models.

The field of olfaction has lacked a standardized, quantitative system of odor representation. Giving researchers access to perceived intensity, and disentangling it from concentration, is a major step toward elevating stimulus control in olfactory research to that of audition and vision.

## Results

Olfactory research has faced significant challenges in precisely controlling odorant concentrations, often leading to discrepancies in reported detection thresholds for the same odorant, sometimes by orders of magnitude (Schmidt and Cain 2010). These discrepancies were often attributable to stimulus variability. To precisely link chemical structure to perceived intensity, we developed a closed-system odor delivery method using gas-sampling bags (Figure 1A). Traditional jar and olfactometer stimulus delivery suffers from two main limitations. First, the participant’s sniff mixes the odor with ambient air, diluting the stimulus and making it difficult to deliver a precise, consistent concentration. Second, odors are typically diluted in solvents, which complicates determination of the air-phase concentration (Jennings et al. 2023) (Jennings et al. 2023). Delivering odors via gas-sampling bags minimizes solvent interactions and prevents dilution from external air. Using this system, we delivered at least seven intensity-spaced concentrations of 62 odorants (more than 434 samples; Figure 1B) to a panel trained to measure intensity using the generalized Labeled Magnitude Scale (gLMS) (Methods). The odors were selected to be diverse in physicochemical and in perceptual space (Figure S2). Validating the reliability of our odor delivery system, our panel mean intensity ratings were stable across replicates (Figure 1C; r(60) = 0.97, RMSE = 3.0) surpassing reliability demonstrated in previous studies (r(18) = RMSE 0.65; (Ma et al. 2021)). Thus, gas-sampling bags enable precise, reproducible odorant delivery, crucial for accurate prediction of odor intensity directly from chemical structure.

**Figure 1.**
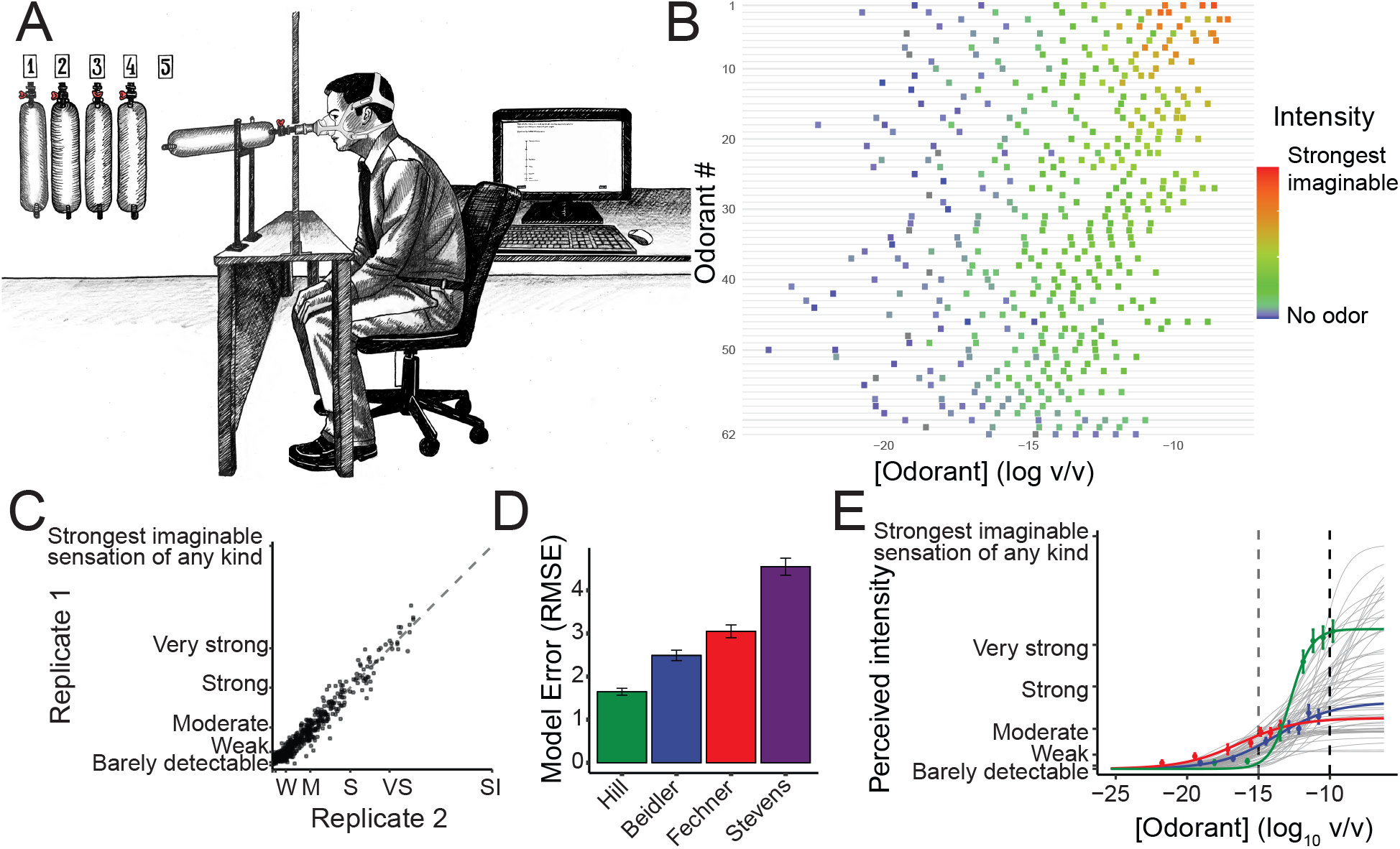
Predicting concentration-intensity relationships of odorants. (A) Experimental setup for collecting intensity ratings using odors in gas-sampling bags. (B) Range of tested concentrations across the sixty-two odorants tested. (C) Human participants’ intensity ratings exhibit high test-retest reliability (panel correlation, r = 0.97) (D) The Hill equation provides the best fit for concentration-intensity data compared to alternative psychophysical models (Beidler, Fechner, Stevens), as indicated by the lowest root mean square error (RMSE). (E) Fitted concentration-intensity functions (Hill equation) for all 62 odorants. The relative intensities of three highlighted odorants—2-heptanone (green), cis-3-hexenyl acetate (blue), and acetophenone (red)—change rank order as concentration increases.

Using this data, we first fitted concentration-intensity curves for each of the 62 odorants. Consistent with previous findings (Chastrette et al. 1998), we confirmed that the Hill equation (Methods, Equation 1) best represents the concentration-intensity relationship in human raters (Figure 1D). These fits demonstrate that the relative intensity ranking between two odorants can depend on concentration (Figure 1E). For example, 2-heptanone (green line) surpasses acetophenone (red line) in perceived intensity as its concentration increases from log (C) = –20 to –15, representing a 105-fold increase. The substantial variation observed in maximum intensity, slope, and inflection point across all 62 odorants further demonstrates that odorant concentration alone is a poor predictor of perceived intensity.

We trained a 5-layer deep neural network (DNN) to predict the fitted values of intensity for each odorant based on their physicochemical features (Figure 2A). Our model performed well on held-out data (RMSE = 7.1, r = 0.87; Figure 2B) and significantly outperformed the same model trained on data where the relationship between intensity and concentration was shuffled (RMSE = 20.4, t(566) = 15.32, p < 0.001). Model accuracy remained consistent across various chemical classes, although prediction errors were most pronounced at maximum intensities (Figure 2C; Figure S3). To further investigate which physicochemical properties drove these predictions, we created a simplified model using only ten select features. These properties included parameters associated with physical transport [molecular weight, vapor pressure, boiling point, and octanol-water partition coefficient (MLOGP) (Mayhew et al. 2022)], a geometric descriptor of molecular shape (alpha-area), and five additional physicochemical features (CATS2D_03_LL, SpMax8_Bh.p., MPC07, MATS7i, H2s) that showed the strongest correlations with the Hill parameters (Fig. 2D, see Methods). The simplified model showed only a ∼1% decrease in predictive performance (RMSE=7.2, r=0.87, Figure 2E), despite using only 10 molecular features compared to 127 in the full model. We explored the correlation between each feature and the three Hill Equation parameters. The maximum intensity (I_max_) was higher for molecules that have a high vapor pressure.

**Figure 2.**
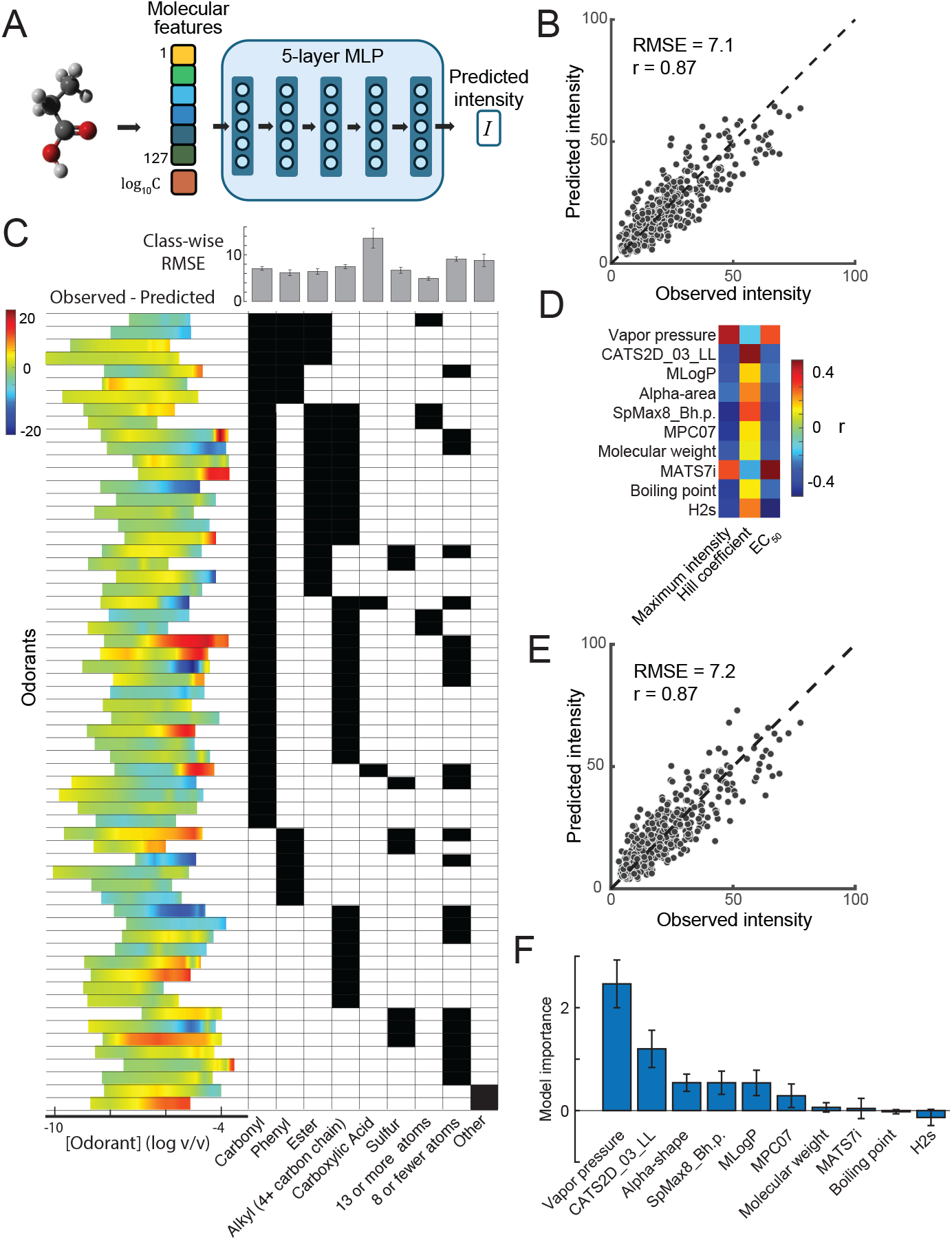
Molecular structure-based prediction of odorant intensity. (A) Schematic of the five-layer neural network trained on 127 physicochemical features, including experimental properties (e.g., boiling point, vapor pressure) and geometric descriptors. (B) Cross-validated model predictions closely matched experimentally measured intensities for held-out odorants (RMSE = 7.1, r = 0.87), significantly outperforming shuffled controls (RMSE = 20.4, p < 0.001). (C) Model accuracy was robust across chemical classes, although prediction errors increased at higher intensities. Colors represent prediction errors (observed – predicted), and grey bars indicate RMSE within each chemical class. (D) Ten molecular features strongly correlated with Hill function parameters of the odorants. (E) A simplified neural network trained on only the 10 selected descriptors achieved nearly equivalent prediction accuracy (RMSE = 7.2, r = 0.87). (F) Key molecular features contributing to predicted odorant intensity include parameters related to odorant transport into the olfactory binding pocket (MLogP, vapor pressure, molecular size (alpha-area)). Feature importance was quantified as the increase in RMSE when each feature was randomly permuted. Error bars show the standard deviation of this RMSE change.

The inflection point (EC_50_) was at lower concentrations for molecules that have a high vapor pressure. The slope of the psychophysical function (Hill coefficient) was the most difficult of the three to predict and was steeper for molecules that have a large number of lipophilic atoms 3-bonds apart (CATS2D_03_LL) (Fig. 2D). To understand how the model uses these features, we randomly shuffled the 10 parameters for the test set of molecules one by one and measured the resulting increase in RMSE (Fig. 2F). The permutation of vapor pressure resulted in a significant RMSE increase (p<10^-5^, bootstrap test, Fig. 2F). Overall, we trained a neural network model that is highly predictive of single molecule intensities and found a set of molecular features relevant to computing perceived intensity.

### Psychophysical curves of individual components enhance mixture odor prediction

Most odors we encounter in everyday life are complex mixtures, yet our understanding of how odorants interact to influence the perceived intensity of mixtures remains incomplete. Historically, simple models (Figure 3A) such as linear addition (ADD) and strongest component (SC) were used to predict odor intensity, though both consistently generate overestimates. Later, geometrical models (Euclidean addition (EUC), vector (VEC), and U-model (U)) incorporated multidimensional relationships between odorants in larger mixtures, improving predictions at the cost of requiring additional empirical parameters. All existing mixture interaction models rely solely on single-concentration intensity measurements for their estimates, have not been systematically compared, and do not incorporate biophysical mechanisms.

**Figure 3.**
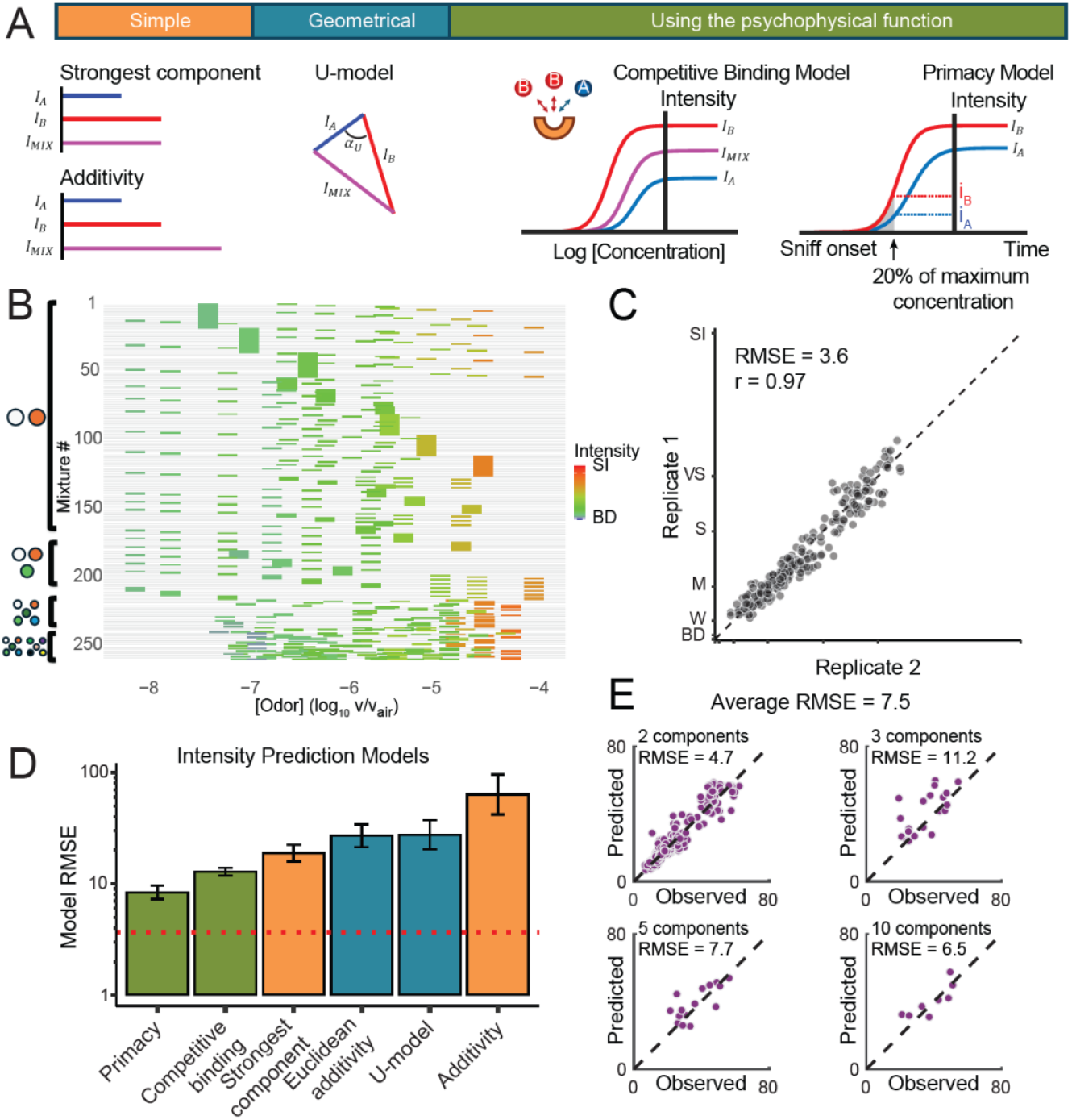
Systematic evaluation of odor mixture intensity models. (A) Overview of historical and biophysical models used for predicting odor mixture intensities. (B) Human panelists rated perceived intensities of odor mixtures composed of 2, 3, 5, or 10 components (7623 ratings total), presented via a standardized delivery system. Each row corresponds to a specific mixture, with colored bars indicating the concentration (x-axis) and intensity (color) of its component odorants when presented alone. (C) Panelist ratings showed high consistency across replicate trials (r=0.97, RMSE=3.62). (D) Models incorporating concentration-intensity functions (green) outperformed traditional models. The red dotted line indicates the test-retest RMSE. (E) A neural network model (same as in Figure 2) accurately predicted odor mixture intensities using component concentrations and physicochemical properties (RMSE=7.5), significantly outperforming a control model trained on shuffled data (RMSE=15.8, p<0.001, bootstrap).

Recently, biophysical models such as Competitive Binding (CB) and Primacy Coding (PRI) have emerged, capturing receptor-level interactions and providing greater biological plausibility (Wilson et al. 2017; Singh et al. 2019). The CB model incorporates concentration-intensity functions into its predictions and here we introduce a modified version of the PRI model, similarly incorporating concentration-intensity functions (Figure 3A; see Methods for details). We hypothesized that incorporating information about each components’ concentration-intensity function would perform better than historical models that only use single-concentration intensity measurements.

To systematically compare these models, trained human panelists (n ≥ 15) evaluated the intensity of odor mixtures containing different numbers of components: 216 binary, 20 tertiary, 16 five-component, and 8 ten-component mixtures. Each mixture was rated twice, resulting in 7623 total data points (Figure 3B). We strategically oversampled mixtures with components at higher intensities, where published models showed the most disagreement. All odorants were delivered via gas-sampling bags, and panelists demonstrated high test-retest reliability (Figure 3C; r(258) = 0.97, RMSE = 3.62) - again surpassing previous studies (r(62) = 0.77; (Ma et al. 2021)).

Supporting our hypothesis, models that incorporated concentration-intensity profiles for individual components outperformed traditional approaches (Figure 3D). Commonly used models, such as EUC and SC, consistently overestimated mixture intensity, as most mixtures were less intense than the strongest individual component. The newly developed primacy model (PRI) achieved the best overall best performance (r(258) = 0.85, p < 0.001, RMSE = 8.6). The primacy model assumes that mixture intensity is determined early in the inhalation cycle, when the odor concentration in the nose is approximately 20% of the ambient concentration (Figure 3A). The intensity of the mixture is then computed as a weighted average using the intensities of individual components at the 20% concentration (Methods, Equation 8).

To predict mixture intensity, the models presented above relied on known intensities or intensity-vs-concentration curves for mixture components. Can quantitative models predict mixture intensity directly from physicochemical properties and concentrations of components? To address this, we trained a DNN with the same architecture as in Figure 2A to directly predict mixture intensity from concentration-weighted physicochemical properties of the components. The DNN was trained end-to-end using both single-molecule response curves and mixture intensity data, with cross-validation to avoid data leakage. This unified approach demonstrated high accuracy in predicting the intensity of held-out mixtures (RMSE = 7.5; Fig. 3E), significantly outperforming a shuffled-label control (RMSE = 15.8, t(316) = 14.1, p < 0.001). These results demonstrate that incorporating both component molecular properties and concentrations enables accurate predictions of mixture intensity.

### Using intensity to identify key odorants in complex aromas

We demonstrate the practical utility of our mixture model by addressing a central challenge in flavor chemistry: identifying the molecules responsible for a food’s aroma. Although foods can emit hundreds of volatile compounds, fewer than 5% meaningfully contribute to their aroma, as most molecules are not present at sufficiently high concentrations to have significant impact (Grosch 2000). Conventional methods rely on headspace solid‐phase microextraction coupled to gas chromatography (HS-SPME-GC) to quantify volatiles, prioritize compounds with an odor activity value (OAV = concentration /detection threshold) > 1 (Dunkel et al. 2014), and then confirm through omission tests (full vs. N−1 blends). This pipeline is slow, expensive, sensitive to variable thresholds (Stevens et al. 1988), assumes intensity is linearly related to concentration (Audouin et al. 2001), and ignores interactions between molecules. Figure 4A highlights this limitation, showing that odorants at the same OAV of 100 span a wide range of intensities, confirming the nonlinear behavior captured by our model.

**Figure 4.**
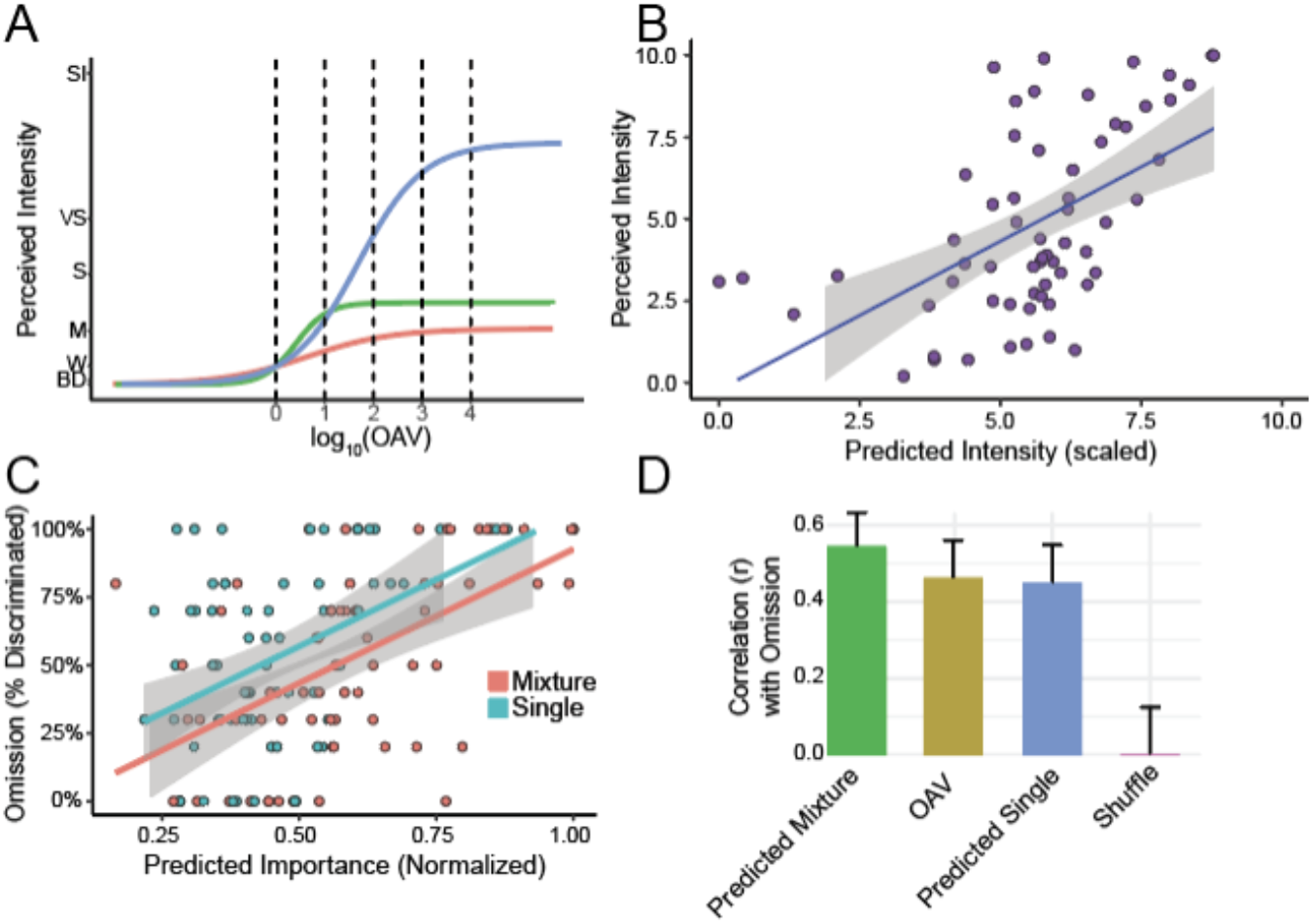
Assessing individual odorants’ contributions to aroma perception. (A) Relationship between odor activity values (OAV) and perceived odor intensities, highlighting substantial nonlinear variation among odorants. (B) Intensities predicted by our single-molecule model strongly correlate with measured intensities of monomolecular components in naturalistic volatile mixtures measured via HS-SPME-GC (r(60) = 0.62, p < 0.001). (C) Single-odorant intensity predictions (as in B) significantly correlate with odorant importance assessed by omission tests (r(65) = 0.45, p < 0.001). Incorporating odorant interactions through neural network (NN) mixture prediction further enhances accuracy (r(65) = 0.55, p < 0.001). (D) The DNN mixture model predictions outperform those based solely on either OAV or the single-molecule intensity model in (B), highlighting the importance of capturing nonlinear interactions when modeling complex aromas.

We propose an alternative to traditional OAV and omission testing by using predictive models to identify key odorants in complex aromas. Drawing on published data (He et al. 2020; C. Liu et al. 2022; R.-S. Liu et al. 2012), we compiled concentrations of 95 odorants across five naturalistic aromas quantified using HS-SPME-GC. In these studies, odorants with OAV > 1 underwent omission testing, comparing the full mixture (N) to a version missing a single component (N-1). Among these odorants, 60 were also evaluated by trained panelists for perceived intensity using GC-Olfactometry (GC-O). First, we externally validated our single-molecule intensity model by predicting the perceived intensity of chemicals rated by GC-O with high accuracy (Figure 4B, r(60) = 0.62, p < 0.001). Next, we evaluated our intensity predictions as an alternative to OAV for identifying key odorants. Although single odorant intensities predicted omission-test outcomes well (r(65) = .45, p < 0.001), our DNN mixture model provided even greater accuracy by simulating intensity reductions between full (N) and omission (N-1) mixtures (r(65) = .55, p < 0.001; Figure 4C). In summary, our intensity-based approach outperformed traditional OAV-based methods in identifying critical odorants (Figure 4D).

## Discussion

In this study, we developed and validated quantitative models to predict perceived odor intensity based on the concentration and physicochemical properties of both individual molecules and complex mixtures.

For individual odorants, we confirmed that a three-parameter Hill function accurately describes the non-linear relationship between concentration and perceived intensity, and related its parameters to molecular descriptors. Extending this model to odor mixtures, we demonstrated that approaches that incorporate the full concentration–intensity profiles of each component outperform traditional additive or “strongest-component” models. Collectively, these advances offer a unified framework for predicting how chemical features translate into perceived intensity, facilitating systematic investigation of odor coding and enabling more precise control of olfactory stimuli in both research and applied contexts.

Without an accurate model to predict the perceived intensity of an odor, neuroscience studies in animal models typically use concentration as a proxy for intensity. While concentration correlates with intensity for a given odor, it fails to predict intensity across odors. For this reason, the field has not yet reached consensus regarding the neural features (e.g., spike rate, latency, ensemble synchrony) underlying perceived intensity. The availability of a reliable intensity metric now makes it possible to design experiments that dissociate concentration from intensity and can adjudicate among neural coding hypotheses such primacy, latency or synchrony models. Additionally, our simplified model uses a small set of molecular transport features, provides greater mechanistic interpretability, and simplifies implementation, thereby broadening its appeal for researchers investigating neural correlates of olfactory perception.

Previous studies have shown that the perceived intensity of odor mixtures does not scale linearly with concentration (Chastrette et al. 1998), and simple additive models often fail to account for mixture interactions like suppression or overshadowing (Cain et al. 1995). Although previous models have been developed to predict odor detection thresholds (Abraham et al. 2012) or odor intensity at fixed concentrations (Keller et al. 2017), these approaches were limited in their ability to generalize across different concentrations or to odor mixtures. Furthermore, limited datasets and challenges in controlling air-phase concentrations have hindered progress (Jennings et al. 2023). Using a large, high-quality dataset of intensity ratings for stimuli with known gas-phase concentrations, we demonstrated that incorporating concentration-intensity functions for individual components substantially improves mixture predictions, outperforming traditional approaches. This is consistent with receptor-level studies showing competitive binding and non-linear responses to mixtures (Singh et al. 2019).

Across sensory modalities, perceived intensity fundamentally shapes stimulus identity. In vision, green light appears brighter than red light even when both deliver identical photon fluxes (Backhaus 1992; Bezold 1873). Similarly, in olfaction, increasing intensity can alter the perceived odor quality of both monomolecular odors (Laing et al. 2003; Gross-Isseroff and Lancet 1988) and complex mixtures (Ravia et al. 2020). However, the field currently lacks a simple, standardized metric for odor intensity, hindering the development of accurate models of odor identity. Current mixture-quality models therefore depend on supplementary intensity ratings from human panelists (Ravia et al. 2020), introducing variability and limiting generalization to novel odors. In food science, this gap means that key aroma compounds are missed because suprathreshold intensity cannot reliably be inferred from detection thresholds. Our mixture model closes this gap by providing an automated, scalable method to predict which odorants dominate perception directly from a gas-chromatography trace.

By converting concentration into a perceptually meaningful scale, our models provide olfactory research with a long-needed analog of decibels or lumens. Just as standardized intensity units have transformed research in vision and audition, these validated intensity models now enable researchers to design stimuli that vary along one perceptual dimension while holding others constant—a fundamental requirement for dissecting neural codes. Odor intensity also governs which components dominate perceptual salience in natural aromas, yet has remained quantitatively intractable. This framework for automated identification of perceptually dominant molecules in complex mixtures has immediate applications in flavor chemistry, fragrance design, and environmental monitoring.

## Methods

### Participants

We recruited 36 panelists (20 female, ages 20-55, mean age = 33.0 ± 11.2 years) from the Philadelphia area. Each testing session used a subset of at least 15 panelists, as panel-averaged intensities are stable at this sample size (Keller et al. 2017). All panelists reported a normal sense of smell and no known health complications that could impair olfaction, such as renal disease, neurodegenerative disease, or congestion at the time of testing. The study was approved by the University of Pennsylvania Institutional Review Board (#818208), and all panelists provided informed consent before enrollment.

### Location and Setup

All experiments were conducted in well-ventilated rooms designed for human olfaction experiments at the Monell Chemical Senses Center. During testing, an opaque curtain obstructed the participant’s view of the experimenter and stimuli (Figure 1A).

### Odor Sampling Bag

Creating a method to deliver known odorant concentrations presents three major challenges: solvent interactions, incompatible solvent mixtures, and dilution from ambient air. The traditional delivery method, an open jar with diluted odorant, fails to control for these factors and often results in poor reliability across participants (Schmidt and Cain 2010). To address these limitations, we developed a controlled odor delivery system using gas-sampling bags (Figure S1). This approach ensures consistent odorant concentrations throughout testing sessions and improves measurement reliability compared to conventional methods. We constructed an odor delivery system using Nalophan gas sampling bags (Miller and McGinley 2008) sealed at both ends with cable ties: one end around a tube fitted with an external septum, the other around an open/close valve. Bags were first evacuated via vacuum through the valve, then filled with dehumidified, carbon-filtered air at a flow rate of 2 LPM for 5 minutes to achieve a volume of 10 L. The liquid odorant was injected through the septum and allowed to equilibrate into the gas phase, a process that took between minutes to hours depending on the odorant’s volume and volatility. After reaching equilibrium, the odorant concentration was uniform throughout the bag. For sampling, we connected the open/close valve to a flexible nose mask equipped with a low-resistance one-way valve. This system allowed natural inhalation of the headspace while preventing ambient air contamination. The mask’s design enabled a tight facial seal and quick attachment to multiple bags in succession. Detailed assembly instructions and a complete materials list are provided in the supplementary material (Table S1 and Document S1).

### Odor Stimuli Selection

We selected 62 odorants (Sigma-Aldrich, >98% purity) to maximize diversity across chemical structure and perceptual qualities. To avoid redundant and trivial stimuli, odorants were chosen to be diverse across two objective criteria: chemical structure (physicochemical features) and perceptual quality (e.g. pleasantness). Compounds were structurally distinct from each other (Figure S2A) and varied in odor character (Figure S2B). A complete list of odorants and their concentrations for both monomolecular and mixture stimuli is provided in Table S2.

### Rating Odor Stimuli Intensity

We trained participants to rate odor intensity using the generalized Labeled Magnitude Scale (gLMS;(Green et al. 1996)) with instructions to focus solely on intensity and disregard pleasantness. Training included ratings of cross-modal verbal examples (e.g., jet engine loudness) and physical weights. Participants were then trained using the bag delivery system and established their baseline response using a blank (clean air) sample. Each testing session began with scale recalibration and included duplicate presentations of both a blank (clean air) and reference stimulus (linalool at -5 log C) to assess rating consistency. Monomolecular odorants were presented at seven or more concentrations, with all stimuli, including mixtures, rated in duplicate. Stimuli were presented in randomized order with 30-second intervals between them. We evaluated participant performance using two criteria: ability to discriminate concentration differences and test-retest reliability within sessions. Participants with poor reliability (rho < 0.6) were excluded from further testing.

### Modeling Monomolecular Concentration-Intensity Functions Psychophysical function for Intensity

Human psychophysical data show that the relationship between an odorant’s concentration and perceived intensity is reminiscent of ligand-receptor binding and can be modeled using the Hill equation (Chastrette et al. 1998):

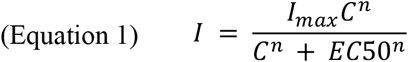

where *I* is the perceived intensity, *C* is the odorant concentration, *I*_*max*_ is the maximum perceivable intensity, *EC*_*50*_ is the concentration at the curve’s inflection point, and *n* is the Hill exponent describing the curve’s steepness. This function produces a sigmoid curve when plotted in log concentration space.

Although other models, such as Stevens’ power law, Beidler’s function, and Fechner’s logarithmic function, have been proposed, Chastrette and colleagues (Chastrette et al. 1998) found that the Hill equation best fit human data. To validate this finding on a larger scale, we fit all four models to our dataset and compared their performance using root-mean-square error (RMSE). The Hill equation consistently yielded the lowest error, and was therefore used throughout this study.

### Analytical Models of Mixture Intensity

Multiple historical models have been proposed to predict odor intensity of a mixture from the perceived intensities (*I*_i_) of its components. We evaluated a range of models grouped into two categories: simple and geometrical. Two additional models using the context of the components’ concentration-intensity relationships were developed specifically for this study. We excluded the vector model from the results as it produced results with imaginary intensity for mixtures with more than 4 components due to the negative cosine term within the square root (see equation 5 below). Mixture intensities were rated by trained panelists using the generalized Labeled Magnitude Scale (gLMS) for binary (N = 216), tertiary (N = 20), five-component (N = 16) and ten-component (N = 8) mixtures. All mixtures were constructed from the 62 odorants that our panelists rated in this study. To maximize contrast between models, we selected component concentrations where model predictions maximally differed, typically above the inflection point of a component’s concentration-intensity curve.

### Evaluation of Model Performance

Model performance was measured using RMSE and Pearson correlation. Since our dataset contained an uneven distribution of mixture sizes, RMSE was computed separately for each mixture type (2-, 3-, 5-, and 10-component mixtures), and then averaged across all groups.

## Simple Models

### Linear Addition (ADD)

Assumes that the perceived intensity of a mixture is the sum of the intensities of its components. For a mixture of *n* odorants, the predicted intensity *I*_*mix*_ can be expressed as:

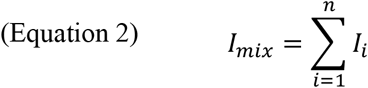

where *I*_*i*_ is the intensity of each odorant when presented alone.

### Strongest Component (SC)

Assumes the mixture’s intensity is determined by the most intense single component:

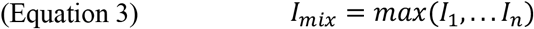

### Geometrical Models

Grounded in Euclidean geometry, these models assume that odorant intensities combine in a multidimensional space and have been shown to provide a more accurate prediction of odor intensity than simpler linear models (Laffort and Dravnieks 1982).

### Euclidean Addition (EUC)

Assumes orthogonal contributions of each component:

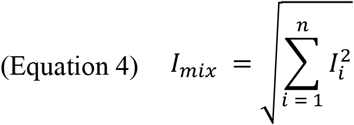

The vector model (VEC) and U model (U), as originally described by (Laffort and Dravnieks 1982), extends the concept of Euclidean addition by incorporating both the magnitudes and angular relationships (cosine similarity) between intensity vectors in a multidimensional odor space. However, each model also requires the estimation of specific parameters for each mixture pair [c*os (α*_*ij*_)] which has previously been derived from the data itself. According to (Laffort and Dravnieks 1982), c*os (α*_*ij*_) of -0.3 will universally work for all mixtures; therefore, this number was used in the analysis.

### Vector Model (VEC)

Extends EUC by incorporating angular relationships (cosine similarity) between odorants:

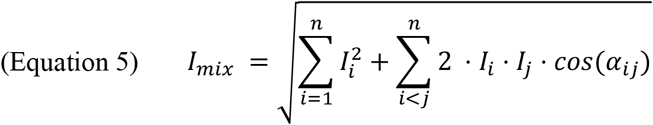

where c*os (α*_*ij*_) is the cosine of the angle between odorant *i* and *j*. Although c*os (α*_*ij*_) can be derived from the data itself, based on prior work (Laffort and Dravnieks 1982), we used a fixed value of –0.3 for all pairwise interactions. Negative cosine values frequently result in imaginary outputs for mixtures with more than three components, leading us to exclude this model from final analyses.

### U Model (VEC)

A more complex model incorporating higher-order interactions among all combinations of odorants:

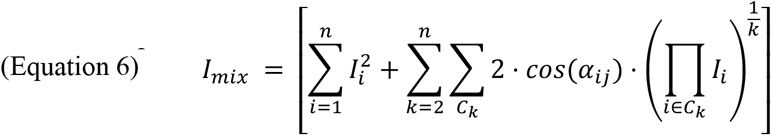

where the first summation represents a linear sum of intensities of individual odorants while the second summation sums the interaction terms for all combinations *C*_*k*_ of *k* odorants, where *C*_*k*_ represents a combination of *k* odorants from the total *n*. The product term accounts for the geometric mean of intensities in each subset.

### Biophysical Models Competitive Binding (CB)

Inspired by receptor–ligand binding dynamics, this model incorporates receptor competition:

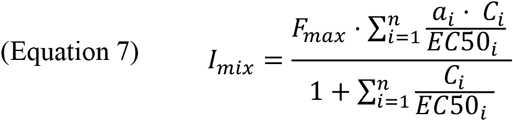

where *a*_*i*_ is the maximal intensity or difference in *I*_*max*_ and *I*_*min*_ for each odorant, *C*_*i*_ is the concentration of each odorant in the mixture, *FC*50_*i*_ is the concentration at half maximal response, and *F*_*max*_ is the global maximum intensity (set to 1). The denominator represents the competition among odorants for binding to the receptor. As multiple odorants compete for the same receptor, the effective concentration available for binding decreases, leading to non-linear responses. While biologically-inspired, it simplifies perception by assuming a single receptor.

### Primacy Coding (PRI)

Based on the theory that early-activated receptor encode odor identity (Wilson et al. 2017), this model estimates mixture intensity from the early part of the concentration-intensity curve, using 20% of the ambient concentration as a proxy for early receptor activation:

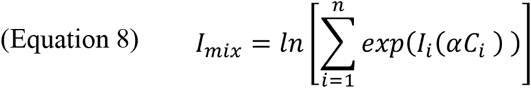

where *I*_*i*_(*αC_i_*) is the intensity of component *i* at 20% (*α* = 0.2) of its maximum (ambient) concentration. Equation 8 is mathematically equivalent to the LogSumExp average of intensities of mixture components taken at 20% of the ambient concentration.

### Deep Learning Models of Intensity

#### Preprocessing of Chemoinformatic Features

For model training, we used 4,885 Dragon molecular descriptors (DDs) generated using Dragon Software (version 6, (Todeschini et al. 2004)). Directly using this high-dimensional feature set would risk overfitting, so we reduced the number of DDs using several steps. First, we discarded DDs which did not display variance for our molecule set (standard deviation < 10-8) or had zero values for most of the molecules. This procedure reduced the number of DDs to 2353. Descriptors with positive values only were log-transformed. Second, to identify descriptors relevant to intensity, we computed correlations between each of the three Hill fitting coefficients for our molecules. For each of the Hill coefficients, we found the 40 most correlated descriptors and included them into our final DD set. We ensured that properties commonly assumed to be predictive of intensity (molecular weight (MW) and Moriguchi octanol-water partition coefficient (MLogP)) were included. Experimental, or secondary estimated boiling points at normal conditions and saturated vapor pressures were gathered from the U.S. Environmental Protection Agencies (EPA) (Estimation Program Interface (EPI) suite, 2024). Additionally, for each molecule, we calculated the bounding shape of 3D molecular structures using MATLAB’s alphashape function. The surface areas of these structures (alpha-areas) were log-transformed and subsequently added to the feature set. After filtering and selection, we retained 127 descriptors for use in the model. The selection process was applied across all molecules, including both train and test data. To assess robustness, we repeated the feature selection within each fold of cross-validation using only training data. The resulting feature sets were highly consistent: 117 of the 127 features appeared in most folds. We observed no meaningful difference in RMSE between uniform and fold-specific feature sets (difference < 1 standard deviation, via bootstrap), so we used the same 127 features across all folds to maintain a consistent model. The same set of parameters was used for predicting both single molecule intensity and the intensity of mixtures.

### Simplified Feature Set for Interpretability

For a reduced model with greater interpretability, we selected a subset of 10 descriptors: five transport-related features (MW, MLOGP, vapor pressure, boiling point, alpha-area) and five descriptors most correlated with Hill coefficients (CATS2D_03_LL, SpMax8_Bh.p., MPC07, MATS7i, H2s).

### Training Data Augmentation

To improve generalization in the single-molecule model, we augmented the training set by upsampling the fitted Hill curves. For each molecule, we sampled the concentration range [10^−10^,10^−3^] at regular intervals, generating 300 additional data points per molecule.

### Single Molecule Intensity Prediction: Data Split

We applied 10-fold cross-validation, stratifying by molecular identity to ensure balanced distribution across folds (Sechidis et al. 2011). Each fold contained unique molecules in the test set, covering all 62 molecules across folds.

### Single Molecule Intensity Prediction: Network Architecture

For each datapoint in the dataset, the pre-processed features were concatenated with the base-10 logarithm of the odorant’s concentration to form an input vector for the model. The neural network (NN) was implemented in MATLAB. The NN consisted of four fully connected layers (N=300), each followed by a batch normalization and a ReLU nonlinearity, with a fully connected (N=1) layer with MSE loss to regress the output with an intensity value.

### Mixture Intensity Prediction: Data Split

For mixture modeling, we used a 90%/10% train-test split with 10-fold cross-validation. All single-molecule training data (62 x 300 data points) were included in the training set. Because the 260 mixtures were constructed from only 24 components, some overlap in single-molecule identities between train and test sets was unavoidable. We assessed the impact of this overlap below.

### Assessing the Impact of Component Overlap

To quantify how overlapping single-molecule identities between train and test sets may influence model performance, we implemented a stricter cross-validation procedure. For each fold, mixtures sharing the same components were grouped together. One group was used as the test set, while mixtures with non-overlapping components were used for training. The single-molecule data included in the training set excluded any molecule present in the corresponding test fold. This “clean split” approach resulted in a test RMSE of **10.5**, higher than the RMSE observed when overlapping components were allowed.

To disentangle the effects of data leakage vs. reduced training size, we matched the number of training samples to the clean split but allowed overlapping molecules. Mixture data (N=260) was randomly partitioned into train and test sets for each fold, matching the train-test distribution of the clean setup. Using this data, we estimated that the impact of test-train set single molecule overlap is 1.0 RMSE units (Figure S4).

### Mixture Intensity Prediction: Network Architecture

To represent a mixture, descriptors for each component were weighted by their relative concentrations (i.e., each molecule’s concentration divided by the sum of all concentrations) and summed. This weighted vector was then concatenated with the log-transformed total mixture concentration to form the final input vector. Alternative vector consolidation strategies were tested but resulted in worse performance. The neural network architecture matched that of the single-molecule intensity prediction model. The network was trained for 80 epochs using MSE loss, stochastic gradient descent (SGD) with momentum, and a 10^-3^ learning rate.

### Mixture Intensity Prediction: Primacy Model

We developed a separate pipeline of mixture prediction by integrating the single molecule NN with a primacy model. The primacy model uses the intensity predictions of single components at 20% of their concentration in the mixture (see Equation 8) and applies this formula to calculate a mixture intensity prediction.

### Estimating Key Odorants in Mixtures

To assess the contributions of individual molecules in complex mixtures, we simulated omission tests using the trained NN. For each of 5 naturalistic odor mixtures (He et al. 2020; C. Liu et al. 2022; R.-S. Liu et al. 2012), the NN predicted the full mixture intensity (*I*_*mix*_). We then modified the NN input by sequentially omitting each molecule from the mixture, generating a predicted intensity for each modified mixture, *I*_*mix*−*i*_ where *i* represents the omitted molecule. For each molecule *i* in the mixture, a simulated omission score was defined as *I*_*mix*_ − *I*_*mix*−*i*_ and compared to experimentally derived omission scores, where such data were available.

As an additional approximation, we also estimated impact using the single-molecule NN by predicting the intensity of each molecule in the mixture in isolation. We then used the single-molecule intensity as an approximation for the omission score (Figure 4C, D).

## Statistical analysis

All statistical analysis was conducted in the statistical software R (R Core Team 2018) in RStudio 2023.06.1. Alpha level was set to 0.05. Data analysis files can be found in the Supplementary Materials.

## Acknowledgements

This research was supported in part by grants from Ajinomoto Co., Inc, and the NIH (T32DC000014, R01DC013339, U19NS112953, R01DC018455, and R01DC017757; F32DC020380 to RP). The authors would like to thank Elena Nikanorova for the illustration in Figure 1 and Cyrille Mascart for help with data.

