## Supplementary Information for "A quantitative framework for predicting odor intensity across molecule and mixtures"

External Files
Table S1. A complete list of materials to make the odor sampling bags.
Table S2. All odorants and their concentrations used in monomolecular and mixture stimuli.
Document S1. Complete Instructions to make the odor sampling bags.

Figures

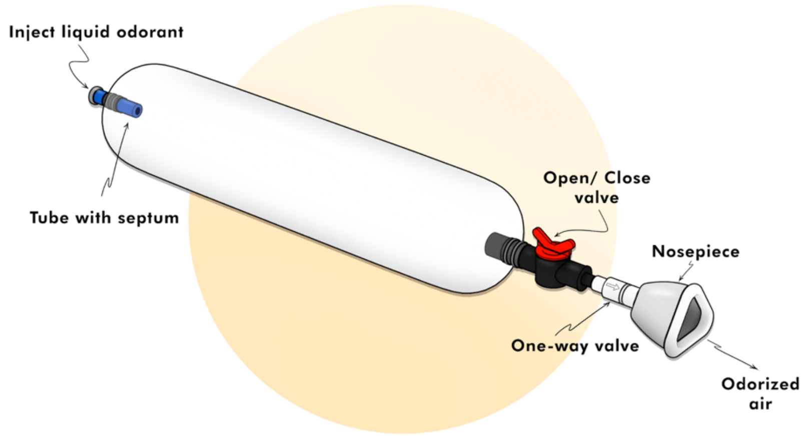


**Figure S1. Schematic of the odor sampling bag**. The bag is made from a tube of thin Nalophan, sealed at one end around a septum for sample injection, and at the other end with an open/close valve to attach to a mask. It maintains a consistent headspace concentration that can be repeatedly sampled without dilution.

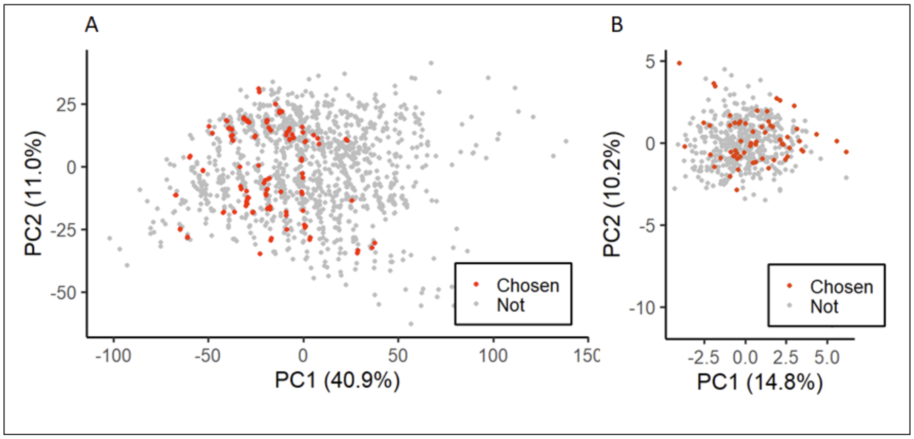


**Figure S2.** Distribution of the selected compounds in (A) physicochemical descriptor space and (B) odor quality descriptor space, visualized using principal component analysis (PCA). Red dots represent compounds that were selected, and gray dots represent a set of ~500 monomolecular odorants.

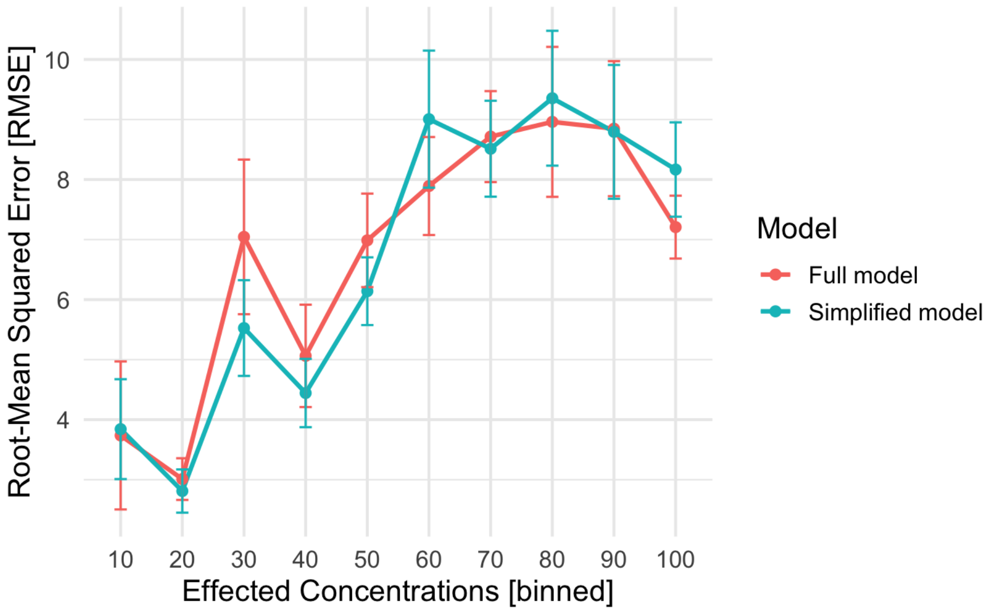


**Figure S3.** Root-mean-squared error (RMSE) is shown for both the full model (red) and simplified model (blue) across binned effective concentrations. Error bars represent standard deviation.

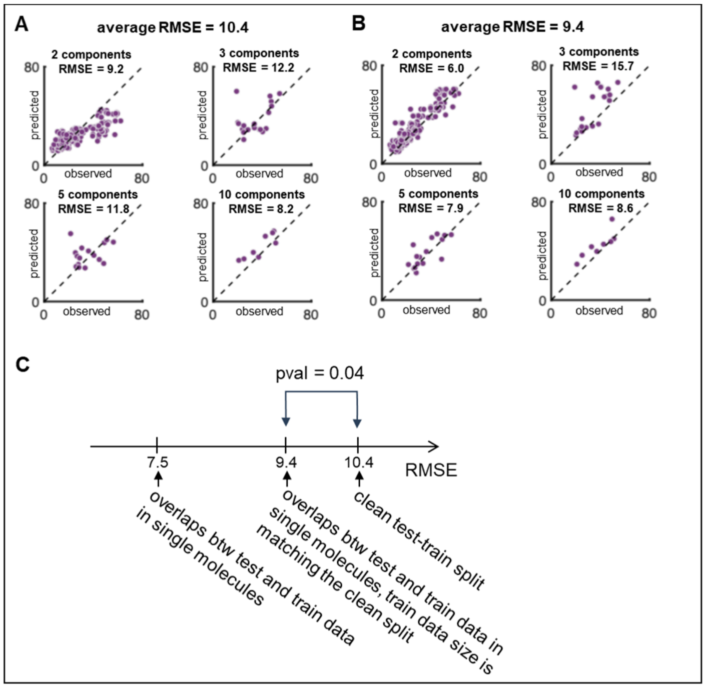


**Figure S4.** Mixture prediction results using a clean split of train and test data. (A) The original mixture dataset included intensity values for mixtures composed of a limited set of monomolecular components. Although the mixture compositions in the training and test sets did not overlap, some individual molecules were present in both sets across folds. Here, we retrained the model using a clean split in which no monomolecular components were shared between train and test data. This yielded a higher RMSE of 10.4. The performance drop compared to the original split (RMSE = 7.5, Figure 3) may be attributed to two factors. First, the removal of overlapping monomolecular components. Second, a reduction in training data size due to this constraint. (B) To isolate the effect of reduced training data size from that of monomolecular overlap, we simulated data scarcity. We trained a model on a dataset that matched the clean split in size and distribution but allowed overlaps in monomolecular components between train and test sets. This yielded an intermediate RMSE of 9.4. (C) Comparing results from (A) and (B), the effect of reduced training data size is estimated as an increase in RMSE of 1.9 units (from 7.5 to 9.4), while the effect of overlapping monomolecular components accounts for an additional increase of 1.0 RMSE units (from 9.4 to 10.4).
